# A virus hosted in malaria-infected blood protects against T cell-mediated inflammatory diseases by impairing DC function in a type I IFN-dependent manner

**DOI:** 10.1101/830687

**Authors:** Ali Hassan, Myriam F. Wlodarczyk, Mehdi Benamar, Emilie Bassot, Anna Salvioni, Sahar Kassem, Antoine Berry, Abdelhadi Saoudi, Nicolas Blanchard

**Affiliations:** Centre de Physiopathologie Toulouse-Purpan (CPTP), Université de Toulouse, Centre National de la Recherche Scientifique (CNRS), Institut National de la Santé et de la Recherche Médicale (Inserm), Université Paul Sabatier (UPS), Toulouse, France

## Abstract

Co-infections shape the host immune status, thereby influencing the development of inflammatory diseases, which can result in detrimental or beneficial effects. For example, co-infections with concurrent *Plasmodium* species can alter malaria clinical evolution and malaria infection itself has the ability to modulate autoimmune reactions but, in both cases, the underlying mechanisms remain ill-defined.

Here, we demonstrate that the protective effects of certain rodent malaria strains on T cell-mediated inflammatory pathologies are due to an RNA virus co-hosted in malaria-parasitized blood. We show that live as well as extracts of blood parasitized by *P. berghei* K173 or *P. yoelii* 17X YM, confer full protection against *Pb* ANKA (*Pb*A)-induced Experimental Cerebral Malaria (ECM) and MOG/CFA-induced experimental autoimmune encephalomyelitis (EAE), and that this is associated with a strong type I IFN signature. We detected the presence of a viral element, the RNA virus Lactate Dehydrogenase-elevating Virus (LDV), in the protective *Plasmodium* stabilates and we established that infection with LDV alone recapitulates the protective effects on ECM and EAE. In ECM, we further show that protection results from an IFN-I-mediated reduction in the abundance of splenic conventional dendritic cell and in their ability to produce the Th1-inducing IL-12p70, leading to a decrease in pathogenic CD4^+^ Th1 responses. In EAE, protection is achieved by IFN-I mediated blunting of IL-12 and IL-23, preventing the differentiation of IFN-γ-, IL-17- and GM-CSF-producing encephalitogenic CD4^+^ T cells.

Thus, our results identify a virus that is co-hosted in several *Plasmodium* stabilates across the community and has major consequences on the host immune system. Moreover, our data emphasize the importance of considering concurrent infections for the understanding of autoimmunity and malaria-associated inflammatory complications.

## Introduction

In extension of the ‘hygiene hypothesis’^1^, the host immune system is ‘conditioned’ by lifelong environmental encounters with micro- and macro-organisms (also known as the multibiome^2^), ranging from commensals to pathogens. In turn, this interplay is expected to influence, positively or negatively, the development and severity of inflammatory disorders including allergy and autoimmune diseases^3^.

With respect to multiple sclerosis (MS), an autoimmune demyelinating disease of the central nervous system, neurotropic virus infections were reported to confer an added risk of developing the disease in adolescents and young adults^4,5^. Although the mechanisms are still debated, it has been hypothesized that virus infections may lead to breakdown of self-tolerance in genetically predisposed individuals, in part by antigenic molecular mimicry and/or by bystander activation of autoreactive T cells *via* upregulation of MHC and costimulatory molecules on dendritic cells (DC)^6^. In experimental autoimmune encephalomyelitis (EAE), a mouse model of MS, viral-specific brain-resident T cells generated in early life following CNS virus infection are indeed able to promote the activity of autoreactive T cells, resulting in brain autoimmune attacks^7^. In contrast to viruses, helminthic parasites are known to play a protective role in MS, most likely because (i) they promote tissue repair and (ii) they induce regulatory T and B cells as well as anti-inflammatory cytokines (e.g. IL-10, TGF-β), which suppress pathogenic autoreactive GM-CSF^+^ CD4^+^ T cell responses (reviewed in^8–10^).

Beside extracellular helminths, intracellular parasites like *Plasmodium,* the malaria-causing parasite, which affects almost half of the world’s population, can also elicit immune modulatory pathways, which could affect autoimmune reactions. Following an asymptomatic liver stage, the parasites reach the blood stream and develop within red blood cells (RBC). Blood stage malaria can be associated with mild to severe clinical symptoms, including anemia, acute respiratory distress syndrome, acidosis, renal failure or cerebral malaria (CM), which is characterized by the sequestration of leukocytes and parasitized RBC (pRBC) in brain microcapillaries, leading to hypoxia and vascular damage. Interestingly, most of the reported modulatory effects of *Plasmodium*, which can affect DC, B and T cells, occur during blood stage. Examples of this modulation on DC include the increased apoptosis of blood-circulating DC^11,12^, an atypical/partial DC maturation profile^13^ and a crippled ability to present antigens to CD4^+^ and CD8^+^ T cells^11,14,15^. The impact of *Plasmodium* on DC may be direct, such as exposure to parasite effectors or byproducts like the heme crystal hemozoin^16^, or indirect, such as the systemic activation by pattern recognition receptors such as Toll-like receptors (TLR), which imprint a ‘refractory’ state on DC^17^, or by type I interferon (IFN-I), which impairs their Th1-promoting property^18^. With regard to T cells, blood stage malaria may causes T cell exhaustion, which can be restored by checkpoint inhibitor therapy^19^. CD4+ T follicular helper (Tfh) normally play a critical role in parasite control during blood stage as they enhance the activation of germinal center B cell responses and enable long-lasting, more efficient humoral immunity^20,21^. Yet during severe malaria, a strong Th1-polarizing environment promotes the development of dysfunctional T-bet^+^ ‘Th1-like’ CD4+ Tfh cells^22,23^, which exhibit poor help activity on B cell responses and lead to B cell apoptosis or differentiation into short-lived plasma cells and atypical memory B cells^24^.

While such immune modulatory processes are thought to partially underlie the poor naturally acquired immunity to malaria observed in endemic areas, they may also have a beneficial impact on the course of autoimmune disorders. More than half a century ago, the incidence of two autoimmune diseases, rheumatoid arthritis and systemic lupus erythematosus, was found to be up to 6-times less frequent in Nigerians than in Europeans, and it was proposed that parasitic infections, in particular malaria, was responsible for alleviating the development of autoimmunity^25^. In accordance, experimental infection with *Plasmodium berghei* (*Pb*) suppressed the spontaneous development of renal disease in a mouse lupic model^26^. Intriguingly, the prevalence and incidence of MS has increased following malaria eradication in Sardinia^27^ and work using rodent-adapted *Plasmodium* strains have reported an overall protective effect of malaria infection on EAE. Infection with *P. chabaudi chabaudi* AS (*Pcc* AS) reduced EAE severity, possibly due to the induction of regulatory CD4^+^ T cells (Treg) and the production of IL-10 and TGF-β^28^. Moreover, transfer of DC incubated with extracts of *Pb* NK65 pRBC ameliorated EAE^29^ although paradoxically, when induced in mice cured from that same parasite, EAE was aggravated^30^. Currently, little is known about the molecular and cellular mechanisms by which *Plasmodium* infection influences CNS autoimmunity.

In addition, beside autoimmune contexts, the clinical evolution of malaria itself is influenced by coinfection with another *Plasmodium* species. In humans, the risk of developing symptomatic malaria seems to be lower in mixed P. *falciparum*/P. *malariae* or P. *falciparum*/P. *vivax* infections^31,32^. In mice, the development of experimental cerebral malaria (ECM), a deadly vascular pathology during which Th1 CD4^+^ T cells promote the sequestration of pathogenic CD8^+^ T cells in the brain vasculature, is inhibited by co-infection with *P. yoelii yoelii* 17X clone 1.1 (*Pyy* 1.1)^33^ and by *Pb* K173^34^. In the former case, replication of the ECM-causing *Pb* ANKA parasites was found to be hampered by mixed infection but in the latter case, there was no effect on parasite growth. Rather, protection was associated with an early production of IFN-γ and IL-10 cytokines at 24 hours post-infection. Yet the exact protective mechanisms remain ill-defined.

Here, in order to elucidate the bases of *Plasmodium*-mediated protection against T cell-mediated inflammatory diseases, we have investigated rodent malaria strains that block ECM and EAE. We show that live blood, as well as blood extracts, parasitized by *Pb* K173 or *Py* 17X YM confer full protection against *Pb* ANKA-induced ECM and MOG/CFA-induced EAE, and that this is associated with a strong IFN-I signature. We report the identification of an RNA virus called Lactate Dehydrogenase-elevating Virus (LDV) in the protective *Plasmodium* stabilates and we find that LDV infection alone recapitulates all the protective effects. In brief, LDV infection triggers a massive IFN-I response, which leads to a decrease in the number and functional capacity of cDC, thereby preventing the provision of signal 3 cytokines that are responsible for the pathogenic polarization of CD4^+^ T cells in ECM and EAE.

## Results

### *Pb* K173 infection does not cause ECM and elicits lower Th1 responses compared to *Pb* ANKA

While the *Pb* ANKA isolate is widely used to induce ECM in C57BL/6 mice, the outcome of *Pb* K173 infection varies. Some studies report its potential to induce ECM^35–37^ while others suggest that it does not cause ECM^38^, or that it even protects from this pathology^34^. We first evaluated the immune responses and clinical outcome following infection with our ‘in-house’ *Pb* ANKA and *Pb* K173 stabilates. We infected mice intravenously (iv) with *Pb* ANKA- or *Pb* K173-infected pRBC, and monitored the development of ECM and circulating parasitemia. Both parasites replicated *in vivo* (**Fig. 1A**) but only *Pb* ANKA led to ECM, characterized by brain edema and early death within 7 to 8 days of infection (**Fig. 1B**). At day 6 post-infection, the absence of ECM development upon *Pb* K173 infection was consistent with a lower number of brain-sequestered CD4^+^ and CD8^+^ T cells compared to *Pb* ANKA (**Fig. 1C**). Moreover, the percentage of brain CD4^+^ (**Fig. 1D**) and CD8^+^ T cells (**Fig. 1E**) producing IFN-γ, an essential cytokine in ECM pathogenesis^39–41^, was significantly decreased upon *Pb* K173 infection compared to *Pb* ANKA. In the spleen, the percentage of CD4^+^ T cells producing IFN-γ upon pRBC restimulation was reduced in *Pb* K173-infected mice (**Fig. 1F**), but no difference was observed in the percentage of IFN-γ^+^ CD8^+^ T cells (**Fig. 1G**). These data suggest that the inability of *Pb* K173 to elicit ECM is related to an impaired differentiation of CD4^+^ Th1 T cells in the spleen and a reduced sequestration of pathogenic CD8^+^ T cells in brain microcapillaries.

**Figure 1.**
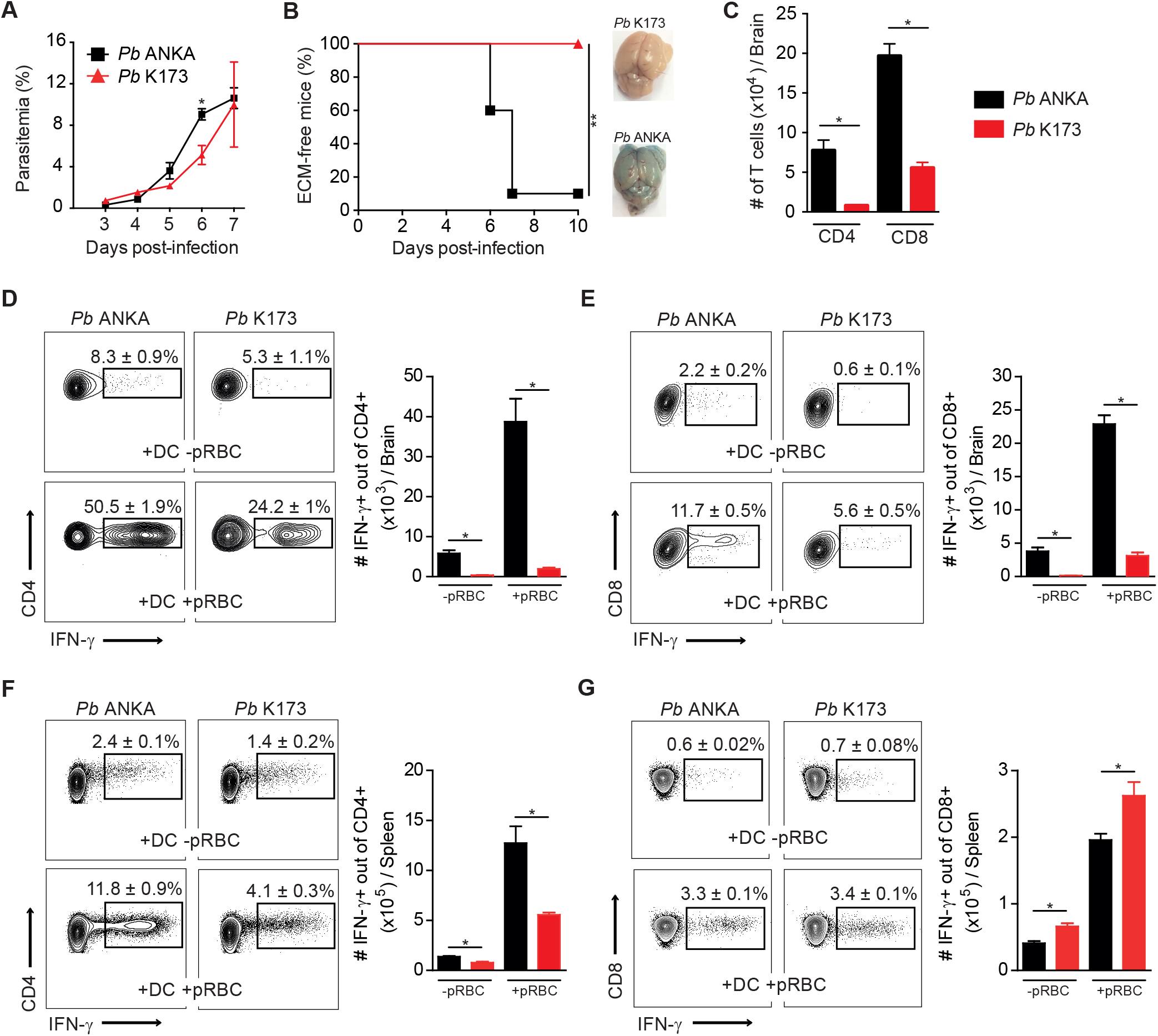
Reduced Th1 responses and absence of ECM pathology following *Pb* K173 infection. C57BL/6 mice infected i.v. by injection of 10^6^ *Pb* ANKA or *Pb* K173 pRBC. Blood circulating parasitemia (A) and ECM development (B) monitored following infection. Brain edema was visualized by Evans blue coloration. (C) Total number of CD4^+^ or CD8^+^ T cells collected from brain at day 6 after infection. (D-G) Cells collected from brain (D, E) and spleen (F, G) at day 6 after infection were restimulated *in vitro* with MutuDC pre-loaded or not with pRBC. IFN-γ production by CD4^+^ T cells (D, F) or CD8^+^ T cells (E, G) detected by intracellular staining. Percentages on the representative dot plots show the mean percentage of IFN-γ^+^ cells out of total CD4^+^ or CD8^+^ T cells +/− SEM. Bar graphs show the absolute numbers +/− SEM. Data are representative of 2 independent experiments. N = 5 mice per group.

### *Pb* K173 infection triggers an early type I IFN response

To investigate the molecular bases behind the lower Th1 responses in *Pb* K173-infected mice, we performed a kinetic analysis of several cytokines in the serum after infection. In accordance with the above findings, bioactive IL-12p70, a hallmark cytokine of Th1 responses, was induced exclusively by *Pb* ANKA but not by *Pb* K173 at day 3 pi (**Fig. 2A**). Instead, *Pb* K173 triggered an early production of IFN-α, CCL4, CCL5, TNF-α, IL-6 and IL-12p40 at day 2 (**Fig. 2A**). In the spleen, *Pb* K173 infection induced a global activation of lymphocytes and DC, as exemplified respectively by the upregulation of CD69 on T cells (**Fig. 2B**), B cells and NK cells (not shown), and of CD86 on DC (**Fig. 2C**). These changes were suggestive of a systemic type I IFN (IFN-I) response^42^. To confirm this hypothesis, we assessed whether they were dependent on IFN-I signaling. We analyzed the cytokine induction and T cell / DC activation in *Pb* K173-infected wild-type *vs. Ifnar1* KO mice. Both the induction of IFN-α, IFN-β, TNF-α and IL-6 genes as well as the systemic activation of splenic T cells and DC were abrogated in *Ifnar1* KO mice (**Fig. 2D, 2E, 2F**). These data indicate that *Pb* K173-but not *Pb* ANKA-parasitized blood triggers a systemic IFN-I response, which correlates with defective Th1 differentiation and with ECM protection.

**Figure 2.**
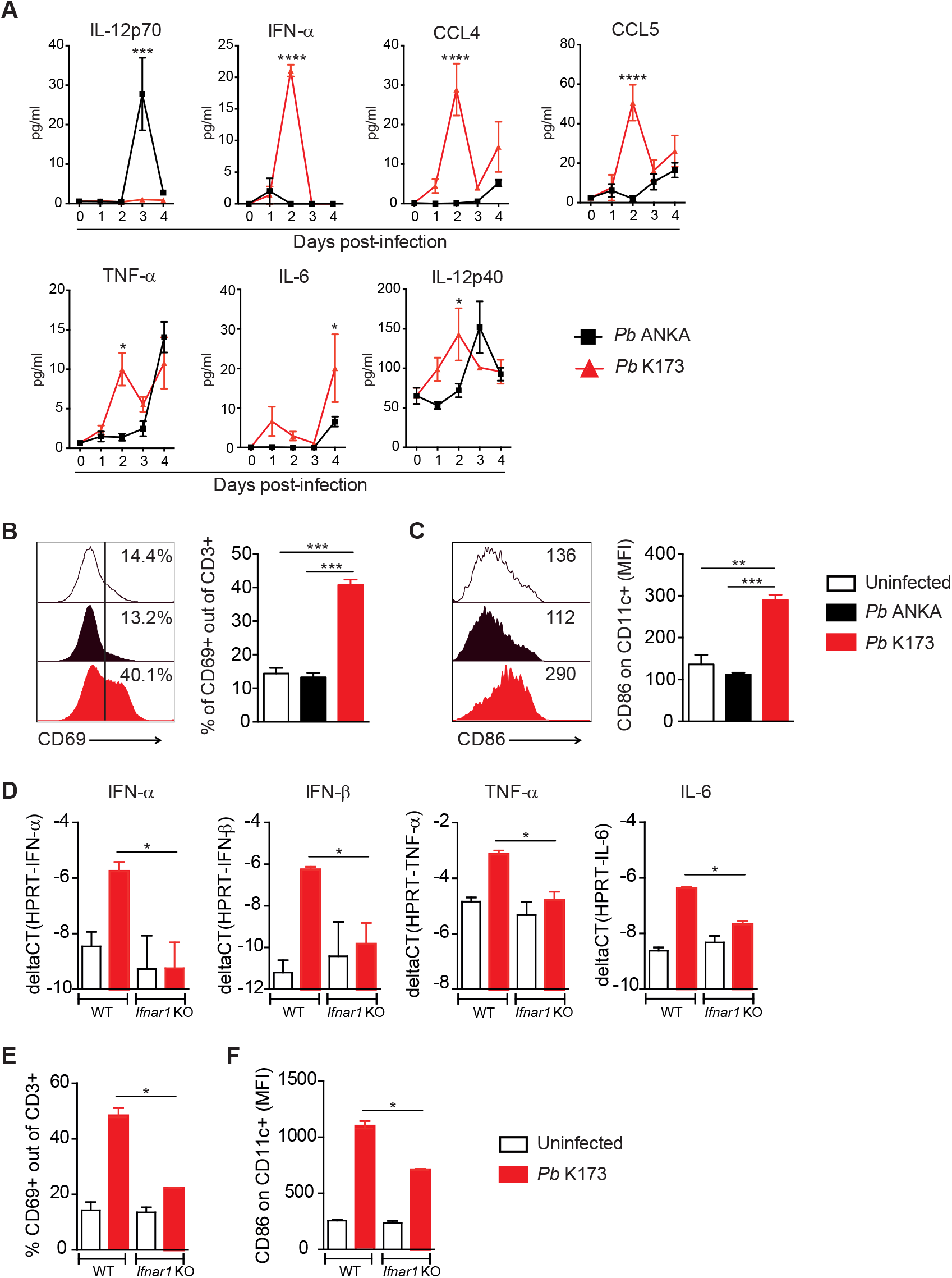
*Pb* K173 but not *Pb* ANKA induces an early systemic type I IFN response. (A) Serum cytokines measured by Luminex assay at different time points after *Pb* ANKA or *Pb* K173 infection. (B, C) Proportion of CD69^+^ out of CD3^+^ T cells (B) and geometric mean fluorescence intensity of CD86 expressed by CD11c^+^ cells (C) in the spleen at day 2 after infection. Bars show the mean +/− SEM. (D-F) WT and *Ifnar1* KO mice were infected with *Pb* K173 and analyzed at day 2 postinfection. (D) Cytokine gene expression in whole spleen analyzed by RT-qPCR. (E, F) Proportion of CD69^+^ out of CD3^+^ T cells (E) and geometric mean fluorescence intensity of CD86 on CD11c^+^ cells (F) in the spleen. Bars show the mean +/− SEM. (A, D, E, F) Data from 1 experiment with 5-7 mice / group. (B, C) Data representative of 4 independents experiments with N=4 mice / group.

### *Plasmodium*-parasitized blood protects against ECM and EAE independently from live parasites

We next investigated if upon *Pb* K173 / *Pb* ANKA co-infection, *Pb* K173 had a ‘dominant’ protective effect on ECM. Co-infected mice were indeed protected from ECM (**Sup. Fig. 1A**) and, consistent with an impaired development of Th1 responses, blood CD4^+^ T cells displayed lower production of IFN-γ (**Sup. Fig. 1B**). It was previously reported that co-infection with distinct clones of *P. yoelii* parasites could inhibit ECM^33^ and that the *P. yoelii* 17X YM (*Py* 17X YM) strain triggers an IFN-I response^43,44^. We thus decided to test whether *Py* 17X YM would phenocopy *Pb* K173. Akin to *Pb* K173, co-infection with *Pb* ANKA and *Py* 17X YM obtained from the MR4/BEI distributor, fully prevented the development of ECM (**Fig. 3A**), and induced T cell and DC activation at day 2 postinoculation (**Fig. 3B, 3C**). These data establish that at least two commonly used rodent malaria strains (*Pb* K173 and *Py* 17X YM) induce a systemic immune activation and protect from ECM.

**Figure 3.**
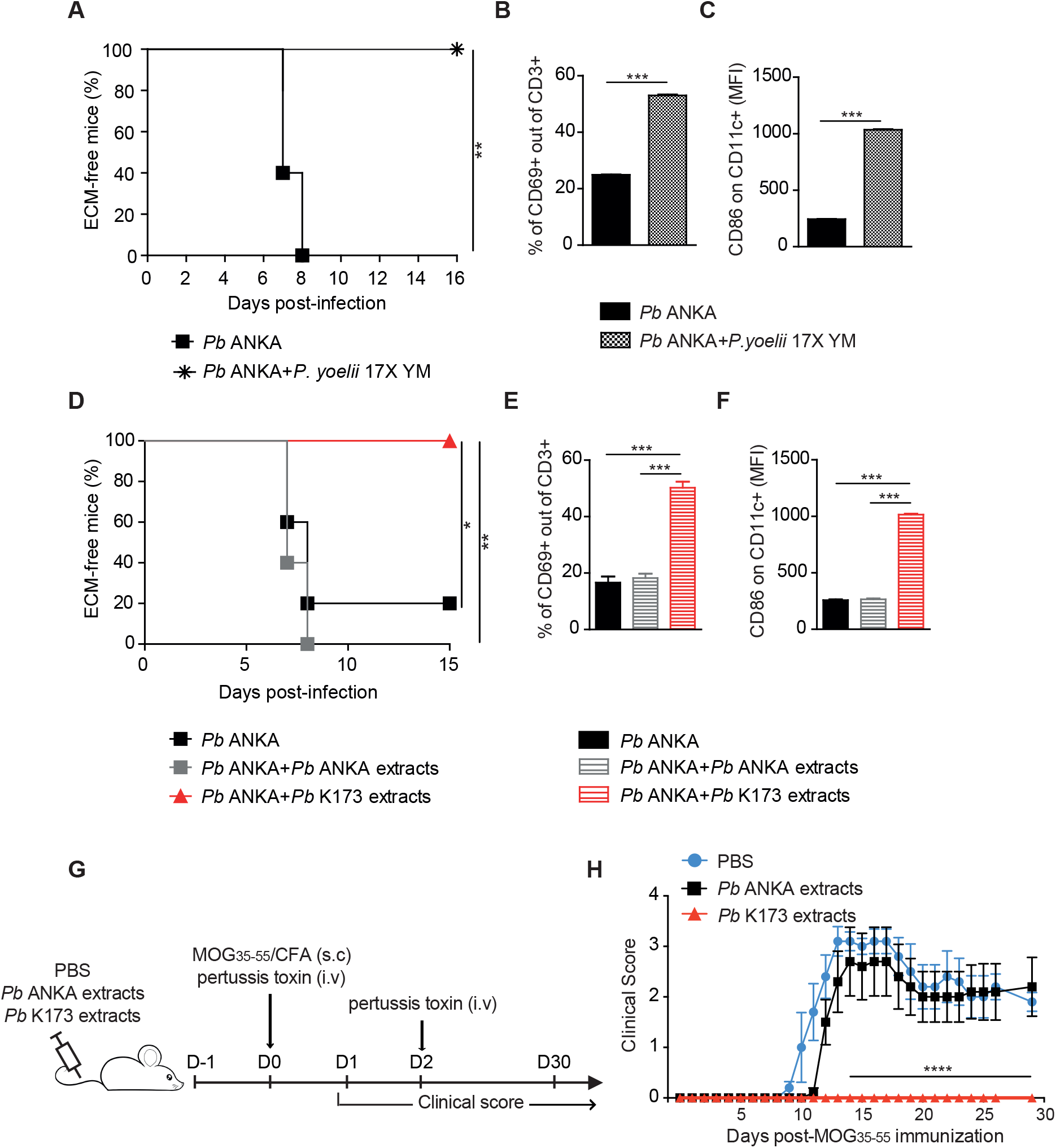
*Plasmodium*-parasitized blood protects against ECM and EAE independently from live parasites. (A) ECM development following single infection with 10^6^ *Pb* ANKA pRBC or mixed infection with 10^6^ *Pb* ANKA pRBC + 10^6^ *P. yoelii* 17X YM pRBC. (B, C) Proportion of CD69^+^ out of CD3^+^ T cells (B) and geometric mean fluorescence intensity of CD86 on CD11c^+^ cells (C) in the spleen at day 2 postinfection. Bars show the mean +/− SEM. (D) ECM development following infection with 10^6^ *Pb* ANKA pRBC administered with or without sonicated extracts of *Pb* ANKA or *Pb* K173. (E, F) Proportion of CD69^+^ out of CD3^+^ T cells (E) and geometric mean fluorescence intensity of CD86 on CD11c^+^ cells (F) in the spleen at day 2 post-infection. Bars show the mean +/− SEM. (G) Experimental protocol. C57BL/6 mice were injected with *Pb* ANKA or *Pb* K173 pRBC sonicated extracts and immunized with MOG_35-55_/CFA to induce EAE. (H) Clinical score monitored up day 30 post-immunization. Dots show the mean +/− SEM. (A, B, C) Data from 2 experiment with N=6 mice per group. (D, E, F) Data representative of 3 independents experiments with N=5 mice / group. (H) Data representative of 2 experiments. N=6 mice / group.

To assess if live parasites were required for protection, we sonicated the *Pb* K173 stabilate, which precluded the parasites from replicating *in vivo* (not shown). Killed *Pb* K173 extracts still conferred full protection against ECM (**Fig. 3D**) and again, protection correlated with a systemic activation of splenic T cells and DC (**Fig. 3E, 3F**). To examine if the protective potential of *Pb* K173 extracts could extend to other T cell-mediated inflammatory diseases, we evaluated their impact on experimental autoimmune encephalomyelitis (EAE), a murine model of multiple sclerosis. EAE was induced by immunization with a MHC II peptide from the myelin oligodendrocyte self-antigen (MOG_35-55_) combined with CFA. Control mice injected with *Pb* ANKA blood extracts 1 day prior to MOG_35-55_/CFA immunization developed a classical EAE disease characterized by a progressive ascendant paralysis but mice that received extracts of *Pb* K173 blood showed no sign of disease (**Fig. 3G, 3H**). This indicates that inhibition of ECM and EAE by *Pb* K173-parasitized blood operates independently from live parasites.

### *Pb* K173 and *Py* 17X YM stabilates contain LDV virus, which, alone, disrupts Th1 responses and protects from ECM

Intriguingly, we noticed that the protective activity of *Pb* K173 blood was contained in the plasma (**Sup. Fig. 2A**) and was disrupted by UV treatment (**Sup. Fig. 2B**), two observations suggestive of the presence of a viral element. To verify this hypothesis, we subjected *Pb* ANKA and *Pb* K173 stabilates to comprehensive PCR Rodent Infectious Agent (PRIA) testing. We found the presence of a virus, the lactate dehydrogenase-elevating virus (LDV), in *Pb* K173 (titer of 3,600 particles per 100 μl plasma) and *Py* 17X YM (titer of 14,600 particles per 100 μl plasma) stabilates, but not in *Pb* ANKA stabilates. LDV is an enveloped, single-stranded positive RNA virus of the family *Arteriviridae,* order *Nidovirales,* known to cause persistent asymptomatic infection in mice, with long-lasting circulating viremia^45^.

This virus has been reported to modulate host responses, thereby either alleviating^46,47^ or exacerbating^48^ immune-mediated diseases. To assess the direct implication of LDV in ECM protection, we decontaminated *Pb* K173 stabilates by Facs-sorting erythrocytes. We verified by PRIA that these new *Pb* K173 pRBC stocks were LDV-free. We also prepared parasite-free LDV-containing plasma by collecting the plasma from mice injected with *Pb* K173 sonicated blood extracts. *Pb* ANKA-infected mice co-injected with LDV-containing plasma were protected from ECM while co-infection with LDV-free *Pb* K173 did no longer protect (**Fig. 4A**). Strikingly, the LDV-free *Pb* K173 stabilate was now causing ECM in the majority of mice (**Fig. 4A**) and these effects were not due to faster parasite growth (**Fig. 4B**). Since we had observed that ECM protection was linked to a reduced Th1 response, we analyzed the activation and production of Th1 cytokines by parasite-specific CD4^+^ T cells at day 6 post-infection. While there was no major difference in the percentages of activated (CD11a^+^ CD49d^+^) splenic CD4^+^ T cells between the groups (**Fig. 4C**), activated CD4^+^ T cells were unable to produce IFN-γ and TNF -α upon restimulation with the I-A^b^-restricted ETRAMP_272-288_ antigenic peptide (**Fig. 4D**). This inhibition was also observed in the total CD4^+^ T cell population upon anti-CD3 stimulation (**Fig. 4E**). The results demonstrate that LDV is necessary and sufficient for ECM protection, most likely by abrogating the Th1 effector activity of CD4^+^ T cells.

**Figure 4.**
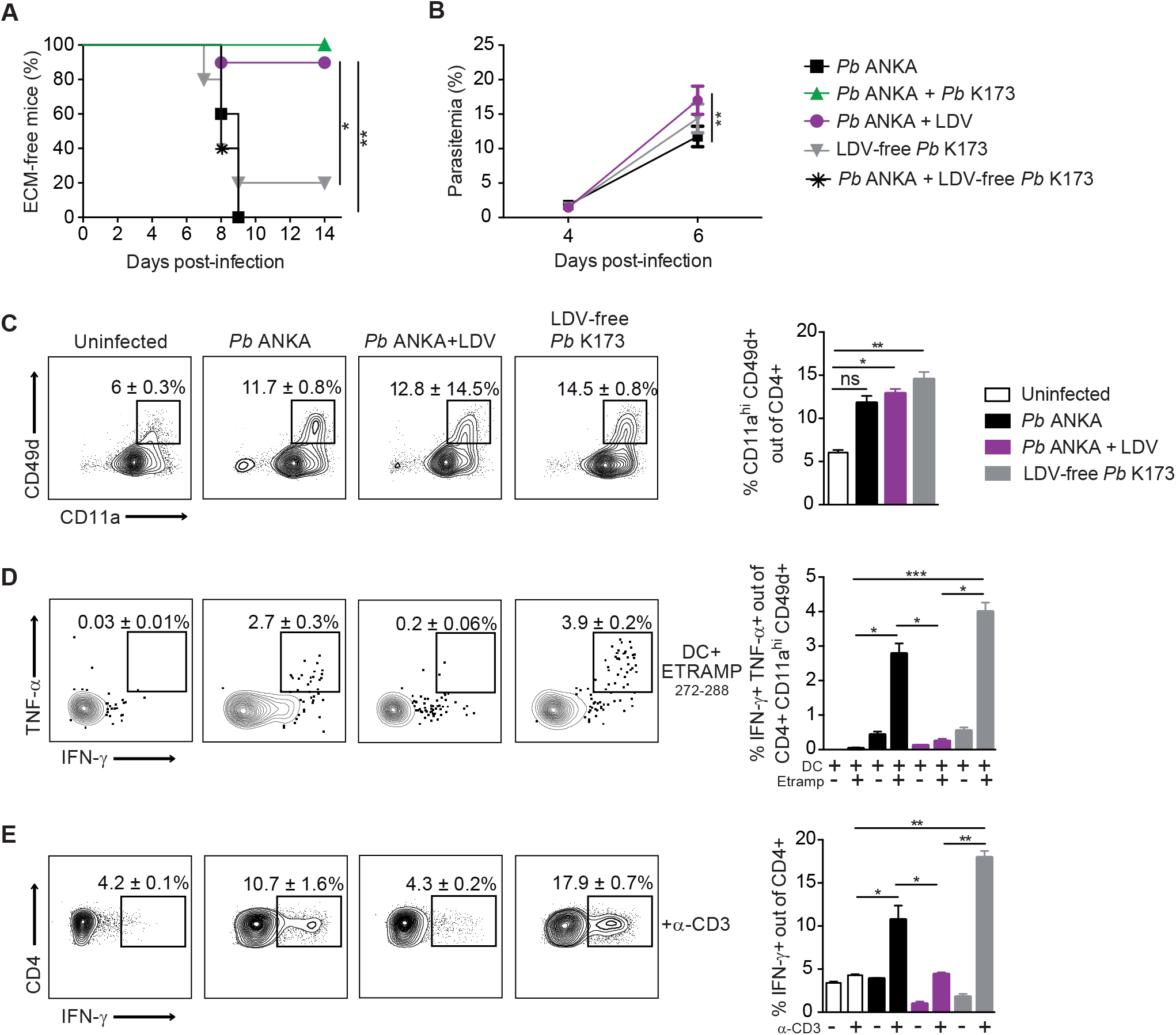
LDV alone, but not LDV-free *Pb* K173, impedes Th1 responses and prevents ECM development. ECM development (A) and blood circulating parasitemia (B) after infection or co-infection of C57BL/6 mice with the indicated inocula: *Pb* ANKA pRBC, *Pb* ANKA + *Pb* K173 pRBC, *Pb* ANKA pRBC + LDV, LDV-free *Pb* K173 pRBC, *Pb* ANKA pRBC + LDV-free *Pb* K173 pRBC. (C) Proportion of activated (CD11a^+^ CD49d^+^) out of spleen CD4^+^ T cells collected 6 days after infection. (D, E) Cytokine production by spleen CD4^+^ T cells restimulated *in vitro* with MutuDC loaded with ETRAMP_272-288_ peptide (D) or with anti-CD3 (E). (D) Proportion of double IFN-γ^+^ TNF-α^+^ cells out of activated CD4^+^ T cells upon ETRAMP272-288 restimulation. (E) Proportion of IFN-γ^+^ cells out of activated CD4^+^ T cells upon anti-CD3 restimulation. Numbers on the dot plots and bar graphs show mean percentages /+-SEM. Data are representative of 2 independents experiments with N=5 mice / group.

### LDV induces IFNAR-dependent depletion and functional impairment of splenic cDC

To study how LDV impedes the Th1 CD4^+^ T cell response, we analyzed splenic conventional DC (cDC), which are primarily responsible for Th1 priming and polarization in this context^49–51^. Two days following *Pb* ANKA / LDV co-infection, we observed a strong reduction in the proportion and numbers of total CD11c^+^ MHCII^+^ cDC, which was not observed in *Ifnar1* KO mice (**Fig. 5A, 5B, 5C**). In proportion, the decrease was more pronounced for the CD8α^+^ cDC1 subset but the CD172a^+^ cDC2 were also affected (**Fig. 5D, 5E, 5F**). In addition, we analyzed the capacity of the remaining cDC to make the pro-Th1 cytokine IL-12p70, which is composed of the IL-12p35 and IL-12p40 subunits. Upon day 2 of infection, *Pb* ANKA induced the upregulation of the IL-12p35 and IL-12p40 genes, which was blocked by LDV co-infection but rescued in *Ifnar1* KO mice (**Fig. 5G, 5H**). Logically, this resulted in the lack of bioactive IL-12p70 in the serum at day 3 post-infection, an effect that was reversed in *Ifnar1* KO mice (**Fig. 5I**). Taken together, our data establish that IFN-I signaling induced by LDV infection causes a quantitative reduction in cDC and a functional impairment in their ability to polarize Th1 CD4^+^ T cell responses.

**Figure 5.**
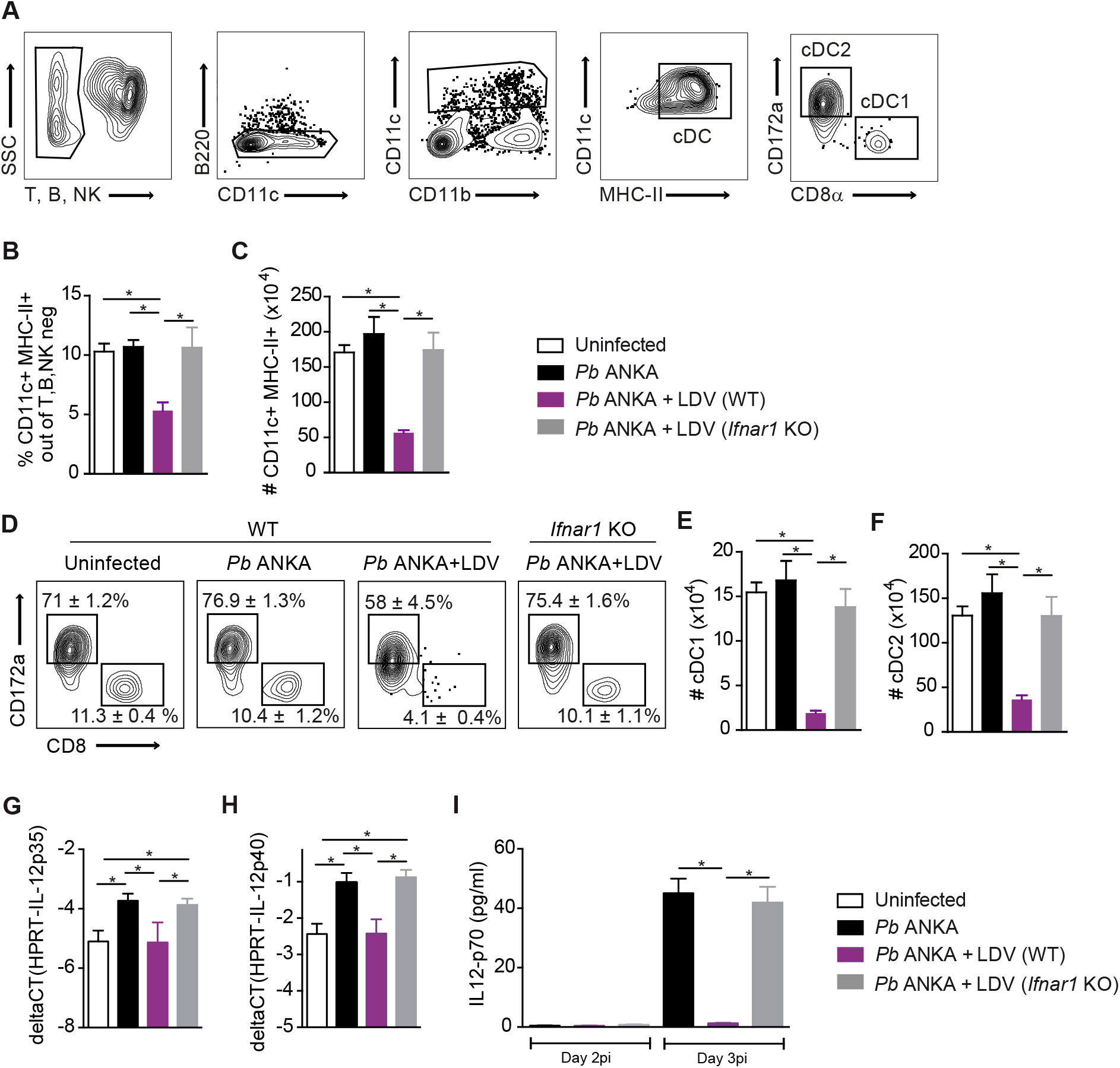
LDV causes an IFNAR-dependent decrease in number and IL-12p70 production of splenic conventional DC. C57BL/6 WT and *Ifnar1* KO mice were infected i.v. with 10^6^ *Pb* ANKA pRBC or 10^6^ *Pb* ANKA pRBC + LDV. (A) Gating strategy used for cDC analysis 2 days after infection. (B, C) Proportion of CD11c^+^ MHCII^+^ cDC out of non-T, non-B, non-NK cells (B) and absolute numbers of cDC (C). (D) Representative dot plots showing the mean percentages +/− SEM of cDC1 and cDC2 subsets. Absolute numbers of splenic cDC1 (E) and cDC2 (F). (G, H) RT-qPCR analysis of IL-12p35 (G) and IL-12p40 (H) gene expression by sorted CD11c^+^ cDC at day 2 after infection. (I) Plasmatic level of IL-12p70 measured by ELISA at day 2 and 3 after infection. (A-H) Data representative of 4 independents experiments with N=5 mice / group. (I) Data representative of 2 independents experiments with N= 5 mice / group.

Type I IFNs can affect multiple cell types including CD4^+^ T cells. Therefore, to determine to which extent the dysfunctional cDC compartment was responsible for LDV-mediated ECM protection, we tested if the transfer of functional cDC could restore disease in *Pb* ANKA / LDV co-infected mice. cDC were isolated from *Pb* ANKA-infected mice and transferred into LDV / *Pb* ANKA co-infected mice on days 3, 4 and 5 (**Fig. 6A**). This procedure restored ECM in only 3 out of 10 transferred mice. We reasoned that even if fully functional at the time of injection, the transferred cDC may immediately be negatively conditioned by IFN-I in the LDV-infected environment. To overcome this effect, we isolated ‘IFN-I-insensitive’ cDC from *Ifnar1* KO *Pb* ANKA-infected mice and transferred them into LDV / *Pb* ANKA co-infected WT mice. This procedure restored ECM in 8 mice out of 10 (**Fig. 6B**), indicating that crippling of cDC by IFN-I signaling is the predominant mechanism that underlies the LDV-mediated protection against ECM.

**Figure 6.**
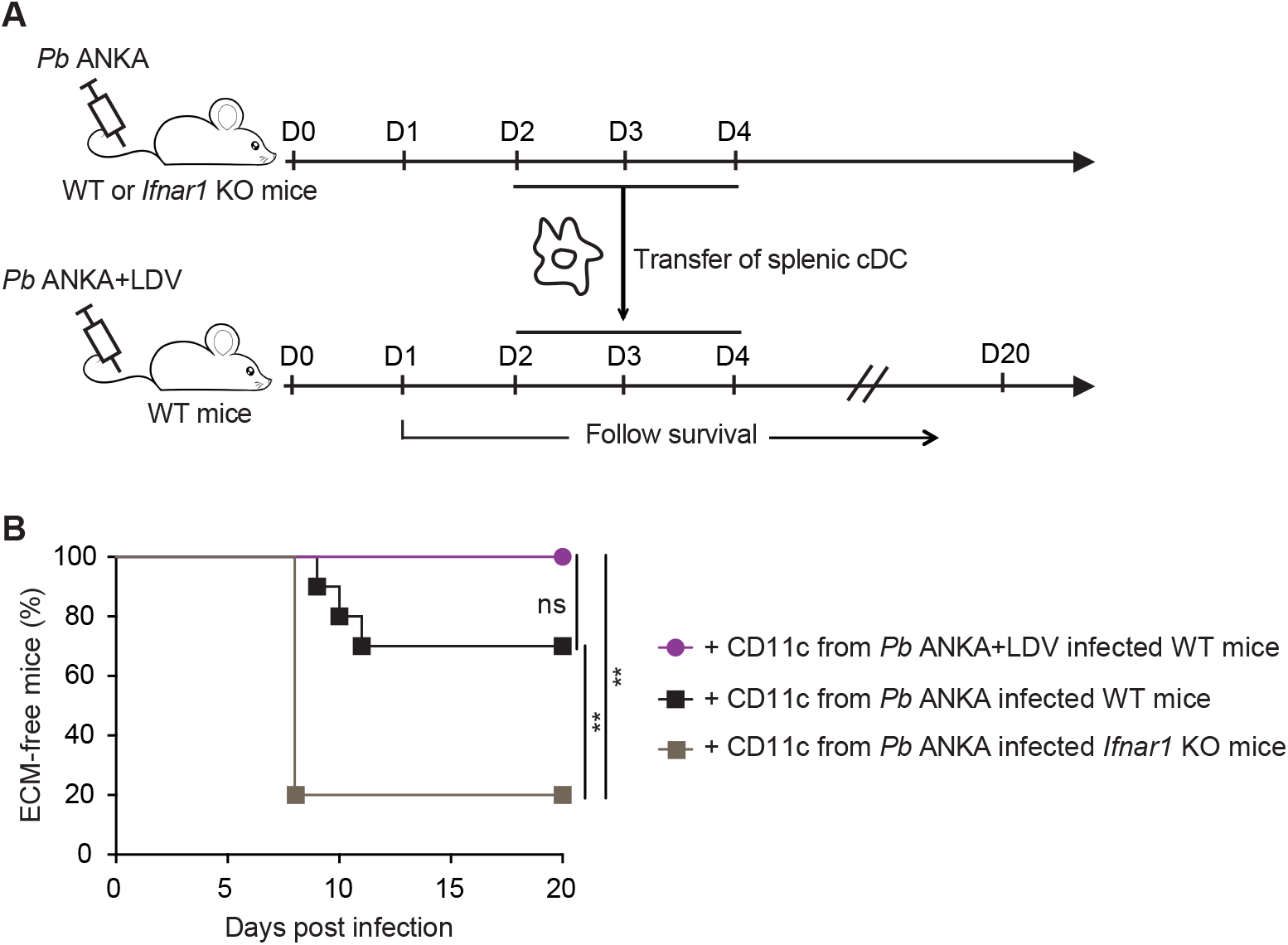
cDC from *Pb* ANKA-infected *Ifnar1* KO mice restore ECM upon transfer into *Pb* ANKA/LDV-co-infected mice. (A) Experimental protocol. WT or *Ifnar1* KO donor mice were i.v. infected with 10^6^ *Pb* ANKA pRBC, with or without LDV. Upon day 2, 3 and 4 of infection, CD11c^+^ DC were magnetically sorted from spleen and transferred into *Pb* ANKA+LDV co-infected WT mice. (B) ECM development. Data pooled from 2 independent experiments with N = 10 mice receiving WT DC and N = 5 mice receiving *Ifnar1* KO DC.

### LDV prevents EAE by disrupting the polarization of pathogenic auto-reactive CD4^+^ T cell responses

Finally, we investigated to which extent the protective mechanisms of *Pb* K173 extracts against EAE were analogous to those described in ECM. We first confirmed that LDV infection alone was protective against EAE. Mice injected with PBS or with sonicated extracts of LDV-free *Pb* K173 prior to MOG_35-55_/CFA immunization developed EAE while mice that received LDV-containing plasma were fully protected (**Fig. 7A, 7B**). Upon day 2 of MOG_35-55_/CFA immunization, the IL-12p40, p35 and p19 subunit genes were induced in cDC isolated from draining lymph nodes (dLN) but this upregulation was blocked by LDV infection in an IFNAR-dependent manner (**Fig. 7C, 7D, 7E**). These findings show that IFN-I signaling blocks the production by cDC of IL-23 (composed of IL-12p40 and p19 subunits), a key cytokine in the polarization of encephalitogenic CD4^+^ T cells^52^. Accordingly, while the proportion of antigen-experienced (CD44^+^) out of CD4^+^ T cells was similar in the spleen and dLN 15 days after MOG_35-55_/CFA immunization (**Sup. Fig. 3A, 3C**), the proportion of autoreactive CD44^+^ CD4^+^ T cells producing IFN-γ, IL-17 and GM-CSF in response to MOG_35-55_ stimulation was lower in LDV-infected mice (**Sup. Fig. 3B, 3D**). This decrease was confirmed by dosing the 3 cytokines in the supernatants of MOG_35-55_-stimulated spleen cells (**Sup. Fig. 3E**). In parallel to the impaired CD4^+^ T cell functional polarization in the spleen and dLN, we observed a major drop in the percentages and numbers of CD4^+^ T cells producing IFN-γ, IL-17 and GM-CSF in the brain (**Fig. 8A**) and spinal cord (**Fig. 8B**), in line with the absence of disease. In conclusion, our data show that LDV-mediated protection against EAE is mediated by an IFN-I signaling-dependent blockade of IL-23-dependent polarization of autoreactive GM-CSF-producing encephalitogenic CD4^+^ T cells.

**Figure 7.**
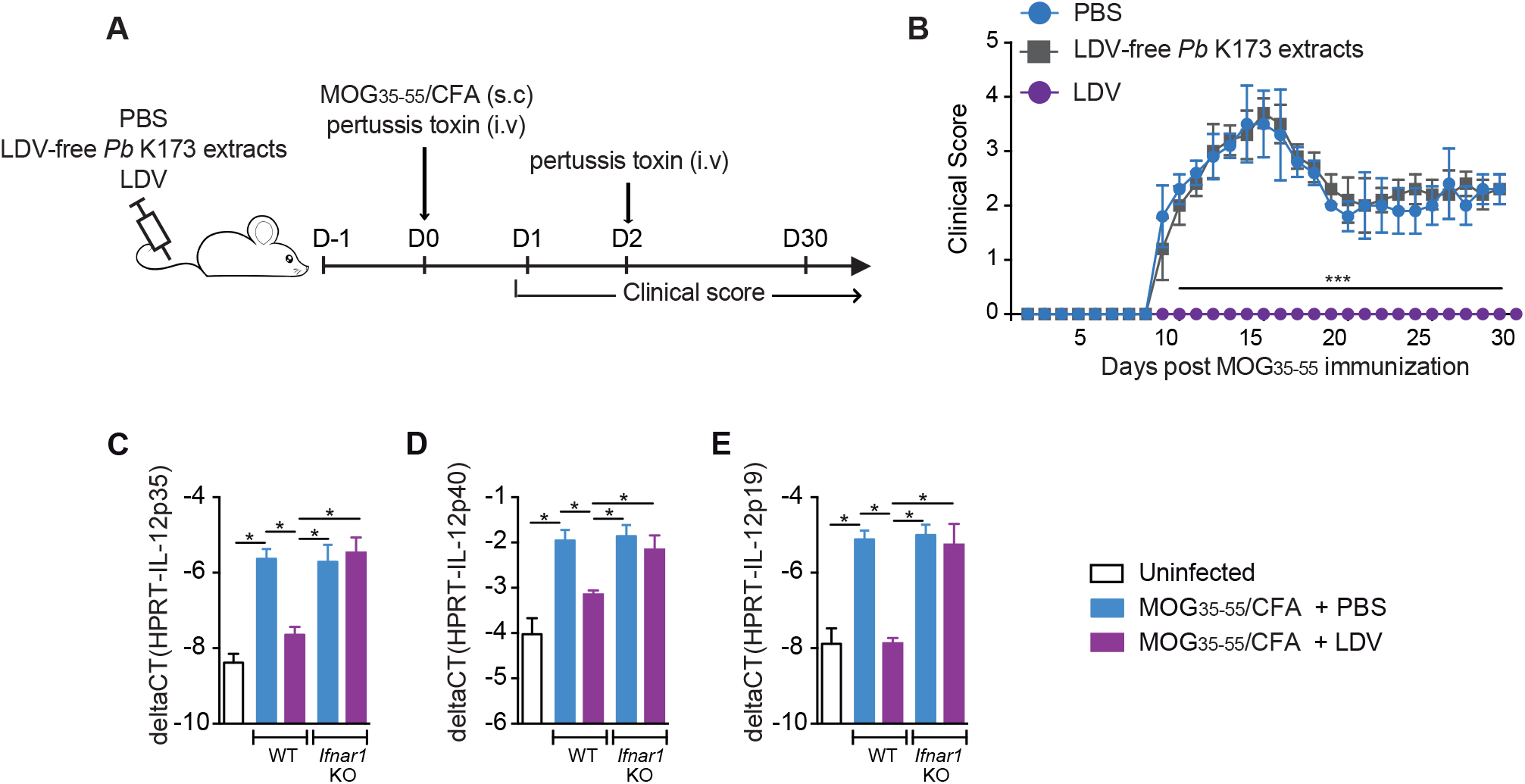
LDV alone, but not LDV-free *Pb* K173, prevents EAE development. (A) Experimental protocol. C57BL/6 mice were injected with PBS, LDV-free *Pb* K173 pRBC sonicated extracts or LDV-containing plasma, and immunized with MOG_35-55_/CFA to induce EAE. (B) Clinical score up to day 30 post-immunization. Dots show the mean +/− SEM. (C, D, E) RT-qPCR analysis of IL-12p35 (C), IL-12p40 (D), and IL-12p19 (E) gene expression on CD11c^+^ DC magnetically sorted from dLN of C57BL/6 WT or *Ifnar1* KO mice, injected with PBS or infected with LDV one day prior to MOG_35-55_/CFA immunization. Analysis at day 1 post-immunization. (B) Data representative of 3 independents experiments with N=5 mice / group. (C, D, E) Data representative of 2 independents experiments with N=5 mice / group.

**Figure 8.**
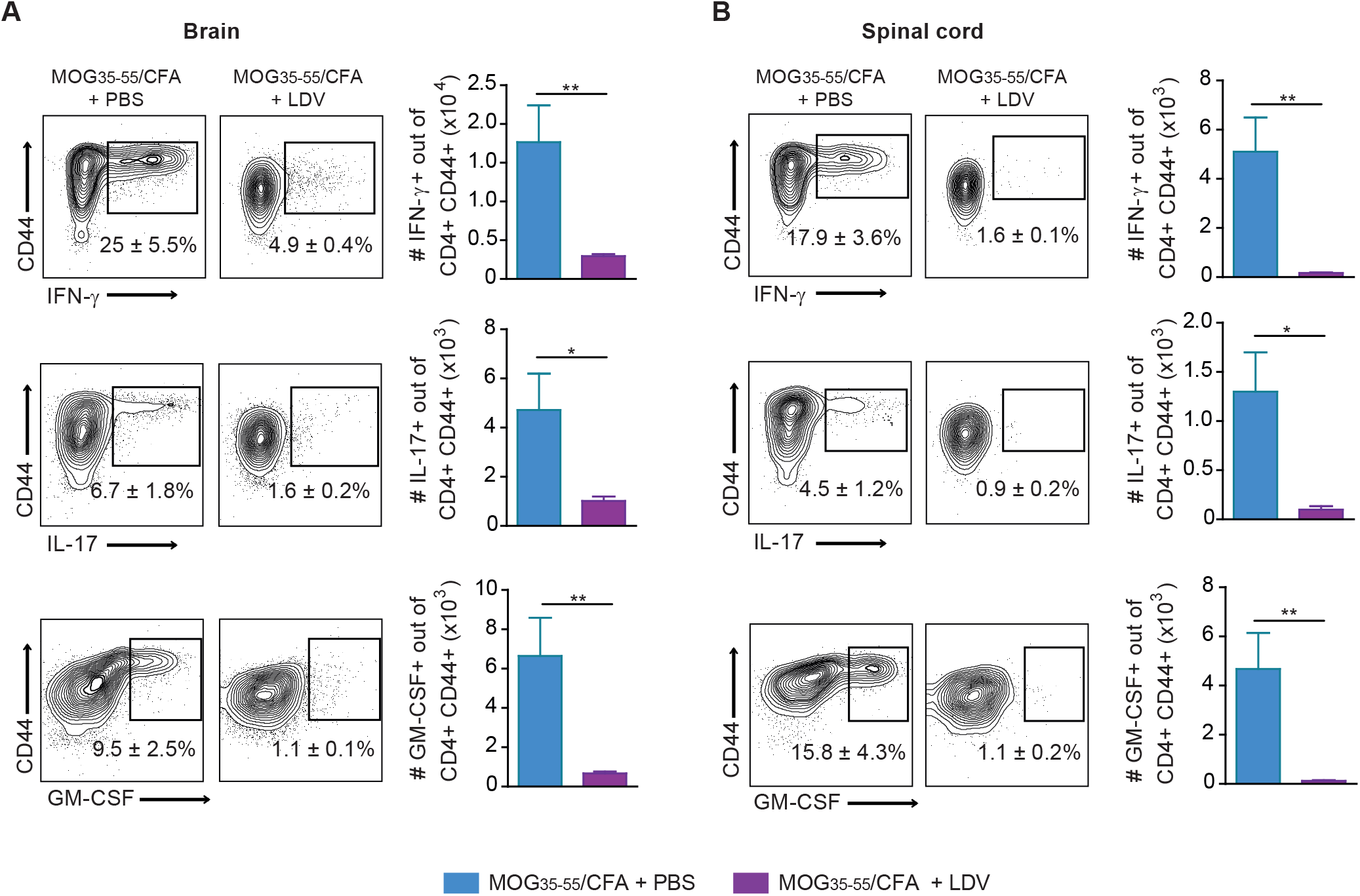
LDV alone protects against EAE by blocking the encephalitogenic CD4^+^ T cell response. C57BL/6 mice were injected with PBS or LDV one day before MOG_35-55_/CFA immunization. Mononuclear cells from CNS and spinal cord harvested 15 days after MOG_35-55_/CFA immunization were restimulated with 100 μg MOG peptide for 24 h. (A, B) Production of IFN-γ, IL-17 and GM-CSF visualized by intracellular staining in CD4^+^ T cells from brain (A) and spinal cord (B). Numbers on the representative dot plots show the mean percentage of cytokine-positive cells out of CD44^+^ CD4^+^ T cells +/− SEM. Histograms show the absolute numbers. Data representative of 2 independent experiments with N = 7 mice / group.

## Discussion

The host immune status is shaped by continuous exposure to commensal as well as potentially pathogenic microorganisms. Depending on whether they exacerbate or alleviate immunopathology, the surrounding microbes may be harmful or beneficial. In this study, we initially sought to address why malaria infection influences the clinical outcome of a concurrent *Plasmodium* infection and autoimmune reactions. To investigate these mechanisms, we took advantage of rodent *Plasmodium* lines that prevent the development of ECM and EAE, two inflammatory diseases mediated by T cells. Unexpectedly, we found that the beneficial effects of the protective *Plasmodium berghei* K173 strain are entirely due to LDV, a non-pathogenic mouse virus, and that this virus is co-hosted in stabilates of other parasite strains (e.g. *P. yoelii* 17X YM) commonly used by the community. We demonstrated that protection against ECM results from an IFN-I signaling-dependent depletion and functional impairment of splenic DC, which become incapable of producing IL-12p70 and polarizing pathogenic Th1 CD4^+^ T cell responses. Analogous to this mechanism, we found that the protection against EAE is linked to an IFN-I-dependent reduction in the production of encephalitogenic IL-23 and GM-CSF cytokines.

This work reiterates the importance of screening products that are injected into animal models for the presence of unsuspected contaminants. In the course of our study, we detected the presence of LDV in stabilates obtained from collaborators (e.g. *Pb* K173, *P. chabaudi*) as well as from a cell line distributor (e.g. *Py* 17X YM from MR4/BEI). LDV was initially discovered as a companion virus of transplantable tumor cell lines more than half a century ago^53^. As it may be co-transmitted along with malaria upon blood stage passages, it was suspected to be present in some *P. chabaudi*, *P. vinckei*, *P. berghei* and *P. yoelii* frozen stabilates^54^. This potential issue seems to have been overlooked thereafter but our data establish that the virus is common in currently used rodent *Plasmodium* stocks. It is possible that some studies on innate immune sensing and/or IFN-I responses performed with LDV-contaminated *Plasmodium* strains may now have to be reinterpreted.

LDV is a mouse-specific virus that causes limited pathology but persists lifelong with detectable circulating viremia^45^. Only one study from 1978 examined the outcome of LDV / malaria coinfection. It was observed that LDV exacerbates *P. yoelii* virulence by increasing parasitemia, leaving open the possibility that the phenotype of the lethal and non-lethal *P. yoelii* 17X substrains may be in part determined by the presence of LDV^55^. Why, in this case, LDV aggravated the pathology remains unclear but one could speculate that with malaria strains that do not cause cerebral malaria, the suppression of Th1 responses is deleterious for parasite control. Concerning autoimmunity, the beneficial role of LDV has been reported in various contexts such as lupus^56^, diabetes^57^ and EAE^46^. In lupus, LDV-mediated protection was correlated with a decrease in IFN-γ production in the serum^58^ but no mechanism was provided to explain the suppression of diabetes and EAE.

Beyond the importance of our findings for the community working on malaria models, our work has fundamental relevance to understand how viruses shape inflammatory diseases. We indeed reveal some of the molecular bases that underlie the protective effects of LDV in two settings involving T cell-mediated immunopathology. In the case of ECM, protection results from the functional paralysis and the strong depletion of splenic cDC, both of which are caused by IFN-I signaling. It was previously shown that LDV induces a rapid systemic IFN-I production by pDC in a TLR7-dependent manner^59^. IFN-I signaling has been reported to play a dual role on the IL-12 pathway. On the one hand, IFN-I drives DC activation and maturation^60^, and production of IL-12^61^, which may augment Th1 CD4 responses. On the other hand, IFN-I can selectively suppress IL-12p70^62^. It is likely that IFNI dosage determines the outcome of IFN-I signaling on Th1 responses^63^. While low levels of IFN-I signaling positively regulate Th1 responses, higher doses of IFN-I, as observed here upon massive viral replication in the first 2 days, rather impede the development of Th1. In addition to the functional alterations of cDC, we reported a major drop in the number of splenic cDC that is dependent on IFN-I signaling. It will be interesting to analyze whether the reduction in cDC is due to apoptosis induced by BH3-only proteins, such as the pro-apoptotic factor Bim^64^.

In the case of EAE, LDV infection prior to MOG/CFA immunization blunted the production of IL-12 and IL-23 by cDC in the dLN, resulting in the absence of IFN-γ, IL-17 and GM-CSF production by MOG-specific CD4^+^ T cells, both in the periphery and in the CNS. It has now become clear that IL-12p35^65^, IFN-γ and IL-17^66^ are dispensable for the development of MOG-induced EAE while the key pathogenic cytokines are GM-CSF produced by CD4^+^ Th cells and IL-23, which may be produced by multiple sources^66,67^. Also, in contrast to ECM pathology in which cDC are required for the priming and polarization of Th1 CD4^+^ T cells^49–51^, cDC are not necessary for the priming of encephalitogenic CD4^+^ Th cells in MOG protein-induced EAE^68^ and mice lacking cDC even display an aggravated disease^69^. Based on this, we propose that LDV-mediated protection against EAE is due to the global effect of IFN-I signaling, which inhibits IL-23 production by several different cell types of the dLN, including cDC, and that this prevents the differentiation of GM-CSF-secreting CD4^+^ T cells. In human DC, IFN-β has been shown to inhibit IL-23p19 production by DC, possibly through a STAT3-dependent induction of the negative regulator SOCS3^70^. The exact transcriptional and/or epigenetic reprogramming that cripple the IL-12 and IL-23 synthesis pathways in response to IFN-I remain to be fully unraveled in this context.

Type I IFN are powerful signals that can modify the functionality of many cell types, including CD4^+^ T cells, which play a major role in the two diseases. For example, in EAE, IFN-I signaling in T cells can directly impair Th17 differentiation^71^. During blood stage malaria, CD4^+^ T cell-intrinsic IFN-I signaling induces T-bet and Blimp-1 expression, thereby promoting IL-10^+^ Tr1 responses in mice^72^ as well as in humans^73^. Therefore, it was possible that CD4^+^ T cells impacted by IFN-I may contribute to protection. Because ECM pathology in LDV-infected animals could be restored by transferring activated, ‘IFN-I-insensitive’ (*Ifnar1* KO) DC from *Pb* ANKA-infected mice, we infer that the role, if any, of non-DC in the protection phenotype should be modest.

There have been other examples of viruses modifying the clinical course of a parasitic disease. A recent striking example is the discovery that a double-stranded RNA virus in *Leishmania* parasites (LRV), which is responsible for the exacerbation of the severity of mucocutaneous leishmaniasis *via* a TLR3/IFN-I-dependent pro-inflammatory response^74,75^. A notable difference is that LRV is an endogenous viral element of the parasite itself while LDV is an exogenous mouse-specific virus that is co-hosted in *Plasmodium*-parasitized blood. Nevertheless, it was observed that the aggravation and metastasis of *Leishmania* lesions were not exclusively associated with LRV and that they could also be enhanced by co-infection with exogenous viruses^76^.

In conclusion, our data emphasize the notion that viral infections, even with viruses that are seemingly innocuous, can have dramatic consequences on a concurrent infectious or autoimmune disease. LDV is a strictly mouse-specific virus that is not transmissible to humans, but the IFN-I-mediated modulatory mechanisms highlighted here are likely to be fully relevant in humans. It is known that constant exposure to an immense variety of commensal and non-commensal microbes profoundly shapes our health. This paradigm supports the importance of actively and systematically characterize environmental microbial agents. From a fundamental standpoint, one could expect exciting avenues of investigations that not only could reveal new languages in the host-pathogen cross-talk but also could suggest novel therapeutic immune modulatory molecules.

## Materials and Methods

### Animals

Animal care and use protocols were carried out under the control of the National Veterinary Services and in accordance with the European regulations (EEC Council Directive, 2010/63/EU, September 2010). Protocols inducing pain were approved under code APAFIS#4318-2016022913333485 v4, by the local Ethical Committee for Animal Experimentation registered by the ‘Comité National de Réflexion Ethique sur l’Expérimentation Animale’ under no. CEEA122. C57BL/6J (B6) mice were purchased from Envigo or Janvier (France) with similar results. *Ifnar1* KO mice with at least 5 backcrosses on the C57BL/6 background were a gift from D. Hudrisier and E. Meunier (IPBS Toulouse). All mice were housed and bred under specific pathogen-free conditions at the ‘Centre Régional d’Exploration Fonctionnelle et de Ressources Expérimentales’ (CREFRE-Inserm UMS006). Mice used in experiments were males aged 8-10 weeks. The number of mice and experimental replicates are indicated in the respective figure legends.

### Parasites and experimental infections

*Plasmodium berghei* ANKA (*Pb* ANKA) and *Plasmodium berghei* Kyberg 173 (*Pb* K173) parasites were gifts from S. Picot (University of Lyon, France) and I. Landau (National Museum of Natural History, Paris, France) respectively. The identity of these parasites was confirmed by whole genome sequencing. *Plasmodium yoelii* subsp. *yoelii*, Strain YM (*Py* 17X YM, MRA-755) was obtained through BEI Resources, NIAID, NIH in 2017, contributed by David Walliker. All *Plasmodium* strains were propagated in C57BL6/J mice. To prepare pRBC stocks, mice were bled in heparin and the proportion of pRBC was evaluated by blood smear. The pRBC concentration was adjusted to 10^7^ pRBC/ml in Alsever’s solution with 10% glycerin. One ml aliquots were stored at −80°C. All infections were done by intravenous inoculation of 10^6^ pRBC. For Evans blue staining, mice were injected i.v. with a solution of 1% Evans blue dissolved in 0.9% NaCl, and brain coloration was examined one hour after dissection. To prepare the sonicated blood extracts, total blood was sonicated using an ultrasonic liquid processor (Branson, Sonifier) under the following parameters: 40 mA, pulse 2, 1 min. Blood was sonicated twice with 1 min rest. For UV treatment, plasma was exposed for 40 min.

### Parasitemia quantification

Parasitemia was measured by blood smear or flow cytometry with similar results. For flow cytometry, 3 μl of tail blood was collected using an EDTA-coated microvette (CB 300 K2E, Sarstedt), and diluted in 500 ml PBS. The diluted blood was labeled with antibodies directed against Ter119 (FITC, TER-119 1/30 Miltenyi Biotec), CD71 (PE, C2 1/300 BD Pharmingen) and CD41-PE-Cy7 (MWReg30, 1/100, Biolegend), fixed and permeabilized with 4% PFA with 0.6% saponin for 10 min, followed by DAPI staining.

### LDV detection and dosage, and decontamination of *Pb* K173 blood

For the initial detection of LDV, *Pb* ANKA and *Pb* K173 stabilates were subjected to a Mouse/Rat Comprehensive Clear Panel for PCR infectious agent testing (PRIA, Charles River). Further detection and quantitation of LDV particles were done with a simple PCR LDV test (Charles River) on mouse plasma.

To free *Pb* K173 blood from the virus, C57BL6/J mice were injected with 10^6^ *Pb* K173 pRBC, 10 days after injection, intracardiac blood was collected with heparin and washed with PBS. RBC and platelets were stained with anti-Ter119 (APC, REA847 1/50 Miltenyi Biotec) and CD41 (PE, MWReg30 1/300 BD Biosciences), and then RBC were separated from platelets by magnetic sorting using anti-PE microbeads (Miltenyi Biotec). The Ter119+ CD41-fraction was stained again with the same anti-Ter119 and CD41 antibodies, and RBC were Facs-sorted with an Aria cell sorter (BD Biosciences). New B6 mice were injected with 10^6^ sorted pRBC. Five days later, a new stock of pRBC was prepared as indicated above. Confirmation for the absence of LDV was performed by PCR (Charles River).

### Induction of EAE and clinical score

The MOG_35-55_ (MEVGWYRSPFSRVVHLYRNGK) peptide was purchased from Polypeptide Laboratories (San Diego, CA, USA). Mice were immunized subcutaneously with 50 μg of MOG_35-55_ peptide emulsified in complete Freund’s adjuvant (CFA, BD Difco, Franklin Lakes, NJ, USA) containing 500 μg of killed *Mycobacterium tuberculosis* (Strain H37a, Difco). Mice were injected intravenously with 200 ng of pertussis toxin (List Biological Laboratories, Campbell, CA, USA) at days 0 and 2 postimmunization. Clinical scores were recorded daily by an experimentator blinded to the experimental groups. Scores were assigned as follows: 0: no sign of disease, 1: loss of tone in the tail, 2: hind limb paresis, 3: hind limb paralysis, 4: tetraplegia, 5: moribund.

### Real-time quantitative PCR for gene expression

Total mRNA was extracted from tissue and cells by standard Trizol-chloroform precipitation. For reverse transcription, the iScript cDNA synthesis kit (Bio-Rad) was used. Quantitative PCR (qPCR) reactions were prepared with LightCycler 480 DNA SYBR Green | Master reaction Mix (Roche Diagnostics). Primers are used at 0.2 μM and presented in Supplementary Table S1. qPCR reactions were run in duplicate on a LightCycler 480 System (Roche diagnostics), and cDNA abundance was normalized to the reference gene *Hprt*. Results are expressed as ΔCt (Hprt minus Gene of interest).

### Isolation of spleen, lymph node and CNS leukocytes

Mice were anesthetized with ketamine/xylazine and perfused intra-cardially with cold PBS. Spleens, lymph nodes, spinal cord and brains were collected in complete RPMI (Gibco) supplemented with 10 % vol/vol FCS (Gibco). Splenocytes were mashed through a 100 μm cell strainer (Falcon). Lymph node cells were mashed in a Potter and filtered through a 100 μm cell strainer (Falcon). Brain and spinal cord were collected separately, homogenized and digested with collagenase D (2.5 mg/ml, Roche), DNAse I (100 μg/ml, Sigma-Aldrich), and TLCK (1 μg/ml, Roche) for 30 min at 37°C. Cells were then washed, suspended in 37% Percoll, and layered on 70% Percoll. After a 20 min centrifugation at 800 g the mononuclear cells were collected from the interface. In both cases, erythrocytes were lyzed using ACK buffer (100 μM EDTA, 160 mM NH4Cl and 10 mM NaHCO3).

### *Ex vivo* T cell restimulation

In the ECM model, 10^6^ splenocytes and 3.10^5^ brain leukocytes were incubated for 5 h at 37°C in the presence of brefeldin A (BfA) with MutuDC, a C57BL/6-derived DC line obtained from H. Acha-Orbea^77^. MutuDC were loaded overnight with pRBC (ratio 1 MutuDC: 10 *Pb* ANKA or *Pb* K173 pRBC), or just incubated with anti-CD3 or 5μM of ETRAMP_272-288_ peptide^51^ during the *in vitro* restimulation. Cells were stained with CD4 (APC-Cy7, GK1.5 1/300 BD Pharmingen), CD8α (BV421, 53-6.7 1/300 BD Biosciences), CD11a (PE, 2D7 1/400 BD Biosciences) and CD49d (BV786, R1-2 1/300 BD Biosciences). Intracellular IFN-γ (APC, XMG1.2 1/200 eBioscience) and TNF-α (Alexa Fluor 700, MP6-XT22 1/300 BD Pharmingen) were detected with intracellular Fixation and Permeabilization Buffer Set (eBioscience) according to the manufacturer’s instructions.

In the EAE model, 6.10^6^ cells from spleen, 3.10^6^ from dLN, 3.10^5^ from brain and 3.10^5^ from spinal cord were incubated for 24 h at 37°C with MutuDC (ratio 1 MutuDC: 10 cells) and 100 μg of MOG_35-55_. BfA was added for the last 5h. Cells were stained with CD4 (BV510, RM4-5 1/200 BD Horizon) and CD44 (BV605, IM7 1/300 BD Horizon). Intracellular IFN-γ (AF700, XMG1.2 1/300 BD Pharmingen), IL-17 (AF700, TC11-18H10 1/200 BD Pharmingen) and GM-CSF (PE, MP1-22E9 1/300 Biolegend) were detected with the same kit as above.

### Flow cytometry analysis of dendritic cells

Spleens and dLN were treated with collagenase D (1 mg/ml, Roche) and DNAse I (100 μg/ml, Sigma-Aldrich), minced into small pieces and incubated at 37°C for 30 to 45 min. Cell preparations were filtered using 100 μM cell strainers. For DC identification, cells were stained for CD3 (PE, 145-2C11 1/300 BD Biosciences), CD19 (PE, 1D3 1/300 BD Biosciences) and NK1.1 (PE, PK136 1/300 BD Biosciences) to exclude T, B and NK cells. A mix containing CD11c (PE-Cy7, N418 1/400 Biolegend), B220 (BV510, RA3-6B2 1/900 BD Horizon), CD11b (PE-CF594, M1/70 1/3000 BD Horizon), MHC-II (AF700, M5/114.15.2 1/400 BD Pharmingen), CD8α (FITC, 53-6.7 1/200 BD Biosciences) and CD172a (APC-Cy7, P84 1/200 Biolegend) was used to characterize the different DC subsets. In some experiments, an anti-XCR1 (PE, REA707 1/200 Miltenyi Biotec) was used to confirm the identity of cDC1. All flow cytometry samples were run on a Fortessa (BD Biosciences) and analyzed using FlowJo software.

### Sorting and transfer of splenic dendritic cells

Spleens were injected with 1 ml of PBS supplemented with Liberase TL (0.125 mg/ml, Roche) and DNAse I (100 μg/ml, Sigma-Aldrich), incubated for 30 min at 37°c, and mashed through a 100-μm cell strainer. RBC were lysed and cells washed in MACS buffer. For DC sorting, cells were incubated 10 min with CD11c microbeads (Miltenyi Biotec), and CD11c^+^ cells were magnetically sorted using according to the manufacturer’s instructions. After sorting, 10^6^ CD11c^+^ cells in 100 μl PBS were transferred to recipient mice by i.v. injection.

### Measurements of soluble cytokiness

Enzyme-linked immunosorbent assay (ELISA) was used to measure cytokines in plasma (50 μl) and culture supernatants (72 hours after restimulation). For IL-12p70, the ELISA MAX^TM^ Deluxe Set (Biolegend) was used according to the manufacturer’s instructions. For the other cytokines, 96-well plates (Nunc Maxi Sorp, Biolegend) were coated for 2 h at 37°C with anti-IFN-γ (AN18), anti-IL-17 (TC11-18H10) or anti-GM-CSF (MP1-22E9). Culture supernatants or standards (Peprotech) were incubated for 2 h at 37°C. The plates were then incubated for 2 h with a secondary biotinylated antibody anti-IFN-γ (XMG1.2), anti-IL-17 (TC11-8H4) and anti-GM-CSF (MP1-31G6), followed by 20 min incubation with streptavidin–phosphatase alkaline (Sigma-Aldrich) at 37°C. Plates were revealed by phosphatase alkaline substrate (Sigma-Aldrich), and absorbance was measured at 450 nm minus 540 nm. Luminex analyses were performed by Genotoul Anexplo platform using the kits EP0X10 and EPX020 (eBioscience), according to the manufacturer’s instructions.

### Statistical Analyses

Statistical evaluation of differences between the experimental groups were determined by using two-way analysis of variance followed by a Bonferroni posttest for clinical monitoring and parasitemia, Kaplan-Meier test for monitoring of ECM-free mice, non-parametric unpaired t test for group *vs.* group comparison, Kruskal-Wallis test for multiple group comparisons. All tests were performed with GraphPad Prism 5.00 (GraphPad Software Inc., San Diego, CA, USA). All data are presented as mean ± SEM.

## Acknowledgments

F. L’Faqihi-Olive, V. Duplan-Eche, A.-L. Iscache, L. de la Fuente for technical assistance at the CPTP-Inserm U1043 flow cytometry facility; R. Balouzat and the zootechnicians at INSERM UMS006-CREFRE mouse facility; S. Ménard, M. Boos and members of the Blanchard and Saoudi teams for technical help and discussions; I. Bernard for setting-up the low severity EAE model, H. Acha-Orbea for the MutuDC, D. Hudrisier and E. Meunier for the B6.*Ifnar1* KO mice, S. Picot for the *Pb* ANKA stabilate, I. Landau for the *Pb* K173 stabilate, BEI Resources, NIAID, NIH for the *Plasmodium yoelii* subsp. *yoelii*, Strain YM stabilate.

This work was supported by ‘Institut National de la Santé et de la Recherche Médicale’, Association pour la Recherche sur la Sclérose en Plaques (to NB and AbS), Human Frontier Science Program Organization (CDA00047/2011 to NB), PIA PARAFRAP Consortium (ANR-11-LABX0024 to NB), PIA ANINFIMIP equipment (ANR-11-EQPX-0003 to NB), French Minister of Education, Research and Technology (PhD fellowship to AH and MB).

## Author Contributions

Conceived and designed the experiments: AH, MFW, MB, SK, AbS, NB. Performed the experiments: AH, MFW, MB, EB, AnS, SK. Analyzed the data: AH, MFW, MB, EB, AnS, SK, AB, AbS, NB. Wrote the paper: AH and NB with help of coauthors.

## Declaration of interests

The authors declare no competing interests.

## Supplementary Figures

**Sup. Fig. 1.**
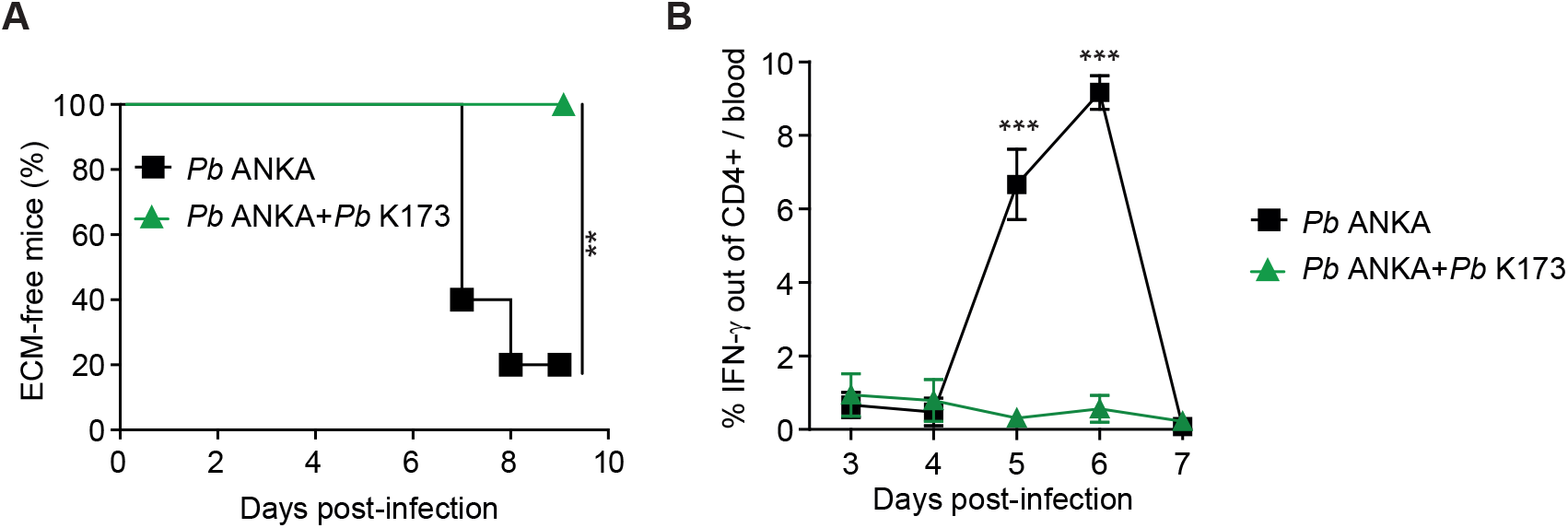
*Pb* K173 co-infection dampens *Pb* ANKA-induced IFN-γ production by CD4^+^ T cells. (A) ECM development in C57BL/6 mice infected with 10^6^ *Pb* ANKA pRBC or 10^6^ *Pb* ANKA pRBC + 10^6^ *Pb* K173 pRBC. IFN-γ expression by CD4^+^ T cells from peripheral blood measured from day 3 to day 7 after infection (B). Data from 1 experiment with N = 5 mice per group. Dots show the mean +/− SEM.

**Sup. Fig. 2.**
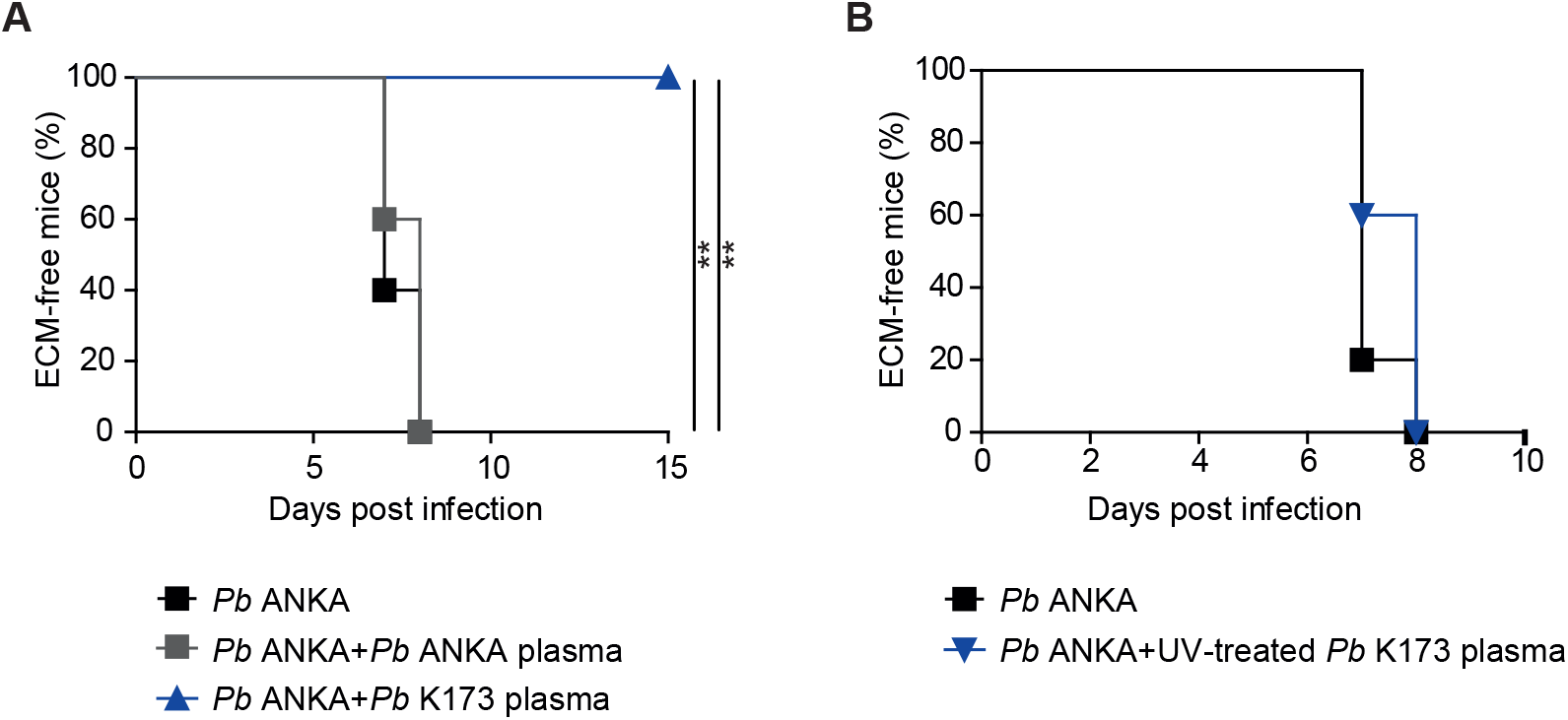
*Pb* K173 protects against ECM independently from the presence of the parasite. (A) ECM development in C57BL/6 mice infected with 10^6^ *Pb* ANKA pRBC and injected with plasma prepared from mice infected either with *Pb* ANKA or *Pb* K173 pRBC. (B) Same as in (A) except that the plasma from *Pb* K173-infected mice was subjected to ultra-violet (UV) radiation before coinjection with *Pb* ANKA pRBC. (A, B) Data from 1 experiment with N = 5 to 7 mice per group.

**Sup. Fig. 3.**
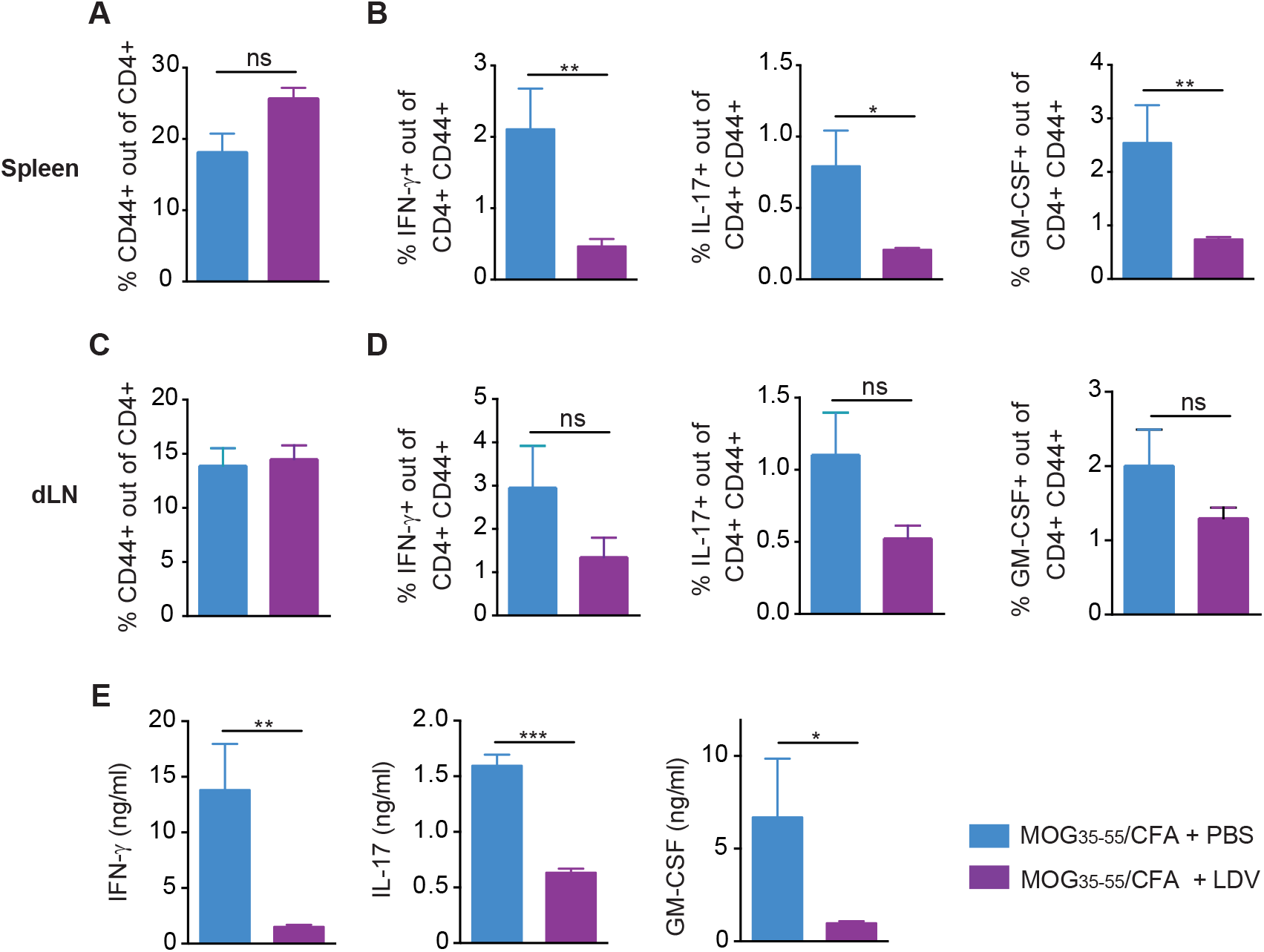
LDV dampens encephalitogenic CD4^+^ T cell differentiation in spleen and draining lymph node following MOG_35-55_/CFA immunization. C57BL/6 mice were injected with PBS or LDV one day before MOG_35-55_/CFA immunization. Mononuclear cells from spleen and dLN harvested 15 days after MOG_35-55_/CFA immunization were restimulated with 100 μg MOG peptide for 24 h. (A, C) Proportion of activated CD44^+^ out of CD4^+^ T cells in spleen (A) and dLN (C). Proportion of CD4^+^ T cells producing IFN-γ, IL-17 or GM-CSF, visualized by intracellular staining, respectively from spleen (B) and dLN (D). (E) Quantification of IFN-γ, IL-17 and GM-CSF cytokines measured by ELISA in the supernatant of MOG_35-55_-stimulated spleen cells. Data representative of 2 independent experiments with N=7 mice / group.

